# Explaining the genetic causality for complex diseases *via* deep association kernel learning

**DOI:** 10.1101/2019.12.17.879866

**Authors:** Feng Bao, Yue Deng, Mulong Du, Zhiquan Ren, Sen Wan, Junyi Xin, Feng Chen, David C. Christiani, Meilin Wang, Qionghai Dai

**Author notes:** These authors contributed equally to this work (Feng Bao, Yue Deng). Corresponding authors: Yue Deng, Meilin Wang and Qionghai Dai.

## Abstract

The genetic effect explains the causality from genetic mutation to the development of complex diseases. Existing genome-wide association study (GWAS) approaches are always built under a linear assumption, restricting their generalization in dissecting complicated causality such as the recessive genetic effect. Therefore, a sophisticated and general GWAS model that can work with different types of genetic effects is highly desired. Here, we introduce a Deep Association Kernel learning (DAK) model to enable automatic causal genotype encoding for GWAS at pathway level. DAK can detect both common and rare variants with complicated genetic effects that existing approaches fail. When applied to real-world GWAS data, our approach discovered potential casual pathways that could be explained by alternative biological studies.

## Introduction

The genome-wide association study (GWAS) is extensively used for uncovering potential causal loci from complex biological phenotypes^1-3^. The classical GWAS models assume that single-locus contributes to the disease independently and the risk increases linearly with the number of minor alleles. These linear models are only powerful in discovering variants with strong and direct associations^4^. As an improvement, pathway-based methods were proposed by taking groups of biologically meaningful genes into consideration^5-7^. For instance, gene-set enrichment methods derive pathway-level statistic scores by combing P-values from single-locus tests ^8-10^; SKAT ^11^ and its variants ^12, 13^ perform association test using kernel regression. However, these existing approaches rely on some pre-assumed genetic models to conduct hand-crafted genotype encoding. Unfortunately, in practice, the genetic effect of complex disease is unknown and can hardly be appropriately modeled in advance. Therefore, a genetic-model-free GWAS approach that can reasonably model the inherent relation between genotype and phenotype is highly needed.

We introduce the Deep Association Kernel learning (DAK) framework to conduct pathway-level GWAS (**Fig. 1**). Our DAK framework incorporates convolutional layers to encode raw SNPs as latent genetic representation. Then, kernel regression layers are connected with these encoded genetic representations to predict the disease status. More importantly, this kernel regression layer allows performing statistical significance tests on the learned genetic representations to uncover the disease-associated pathways. Both the convolutional and kernel regression layers are trained jointly using multiple-instance loss in an end-to-end manner. Therefore, DAK relies on no pre-assumed genetic model and can learn all model parameters in a pure data-driven manner.

**Figure 1.**
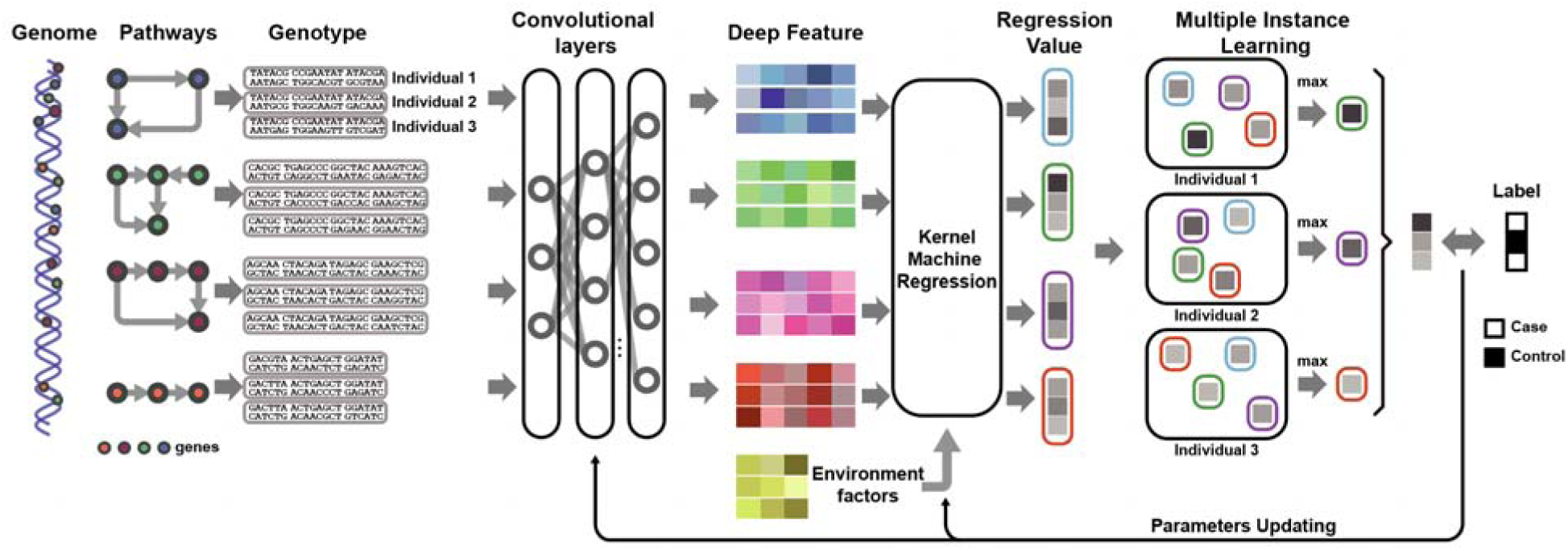
The framework of DAK. SNPs are grouped into pathway-level gene set and coded into one-hot format. Convolutional layers are employed to encode causal loci into deep features. Kernel machine regression is incorporated to enable statistical tests of association via SKAT framework. Multiple instance learning selects the most suspicious pathway at individual level. Parameters of the whole framework are optimized in an end-to-end manner through backpropagation.

We compared DAK with seven representative gene/pathway based methods: classical statistic method (Burden test)^14^, enrichment methods (GATES, HYST and aSPU)^9, 15, 16^ and kernel methods (SKAT and SKAT-o)^11, 12^. DAK is the only approach that consistently performs well under a wide range of genetic models including additive, multiplicative, dominant, recessive and heterozygous effects. We further applied our method to four disease datasets including gastric cancer, colorectal cancer, lung cancer and psychiatric disorder.

## Results

### Deep association kernel learning

We introduced deep association kernel (DAK) learning to achieve the detection of complex associations and enhance the interpretability of GWAS (**Fig. 1** and **Methods**). Here, alleles are coded in the one-hot representations to enable flexible modeling of genotype effects for each locus. Variants in the same biological pathway are grouped together and the combinational effects of multiple SNPs within a pathway are considered at the same time. Then, pathway-level features are extracted by convolutional layers (**Supplementary Fig. 1**), followed by a kernel regression layer to derive the statistical significance (**Supplementary Fig. 2**). To allow learning from labels at individual level, the whole framework is trained with a multiple instance loss in an end-to-end manner. Finally, the variance tests used in SKAT are performed on the learned kernel matrix to derive statistical P-values (**Supplementary Figs. 3** and **4**).

**Figure 2.**
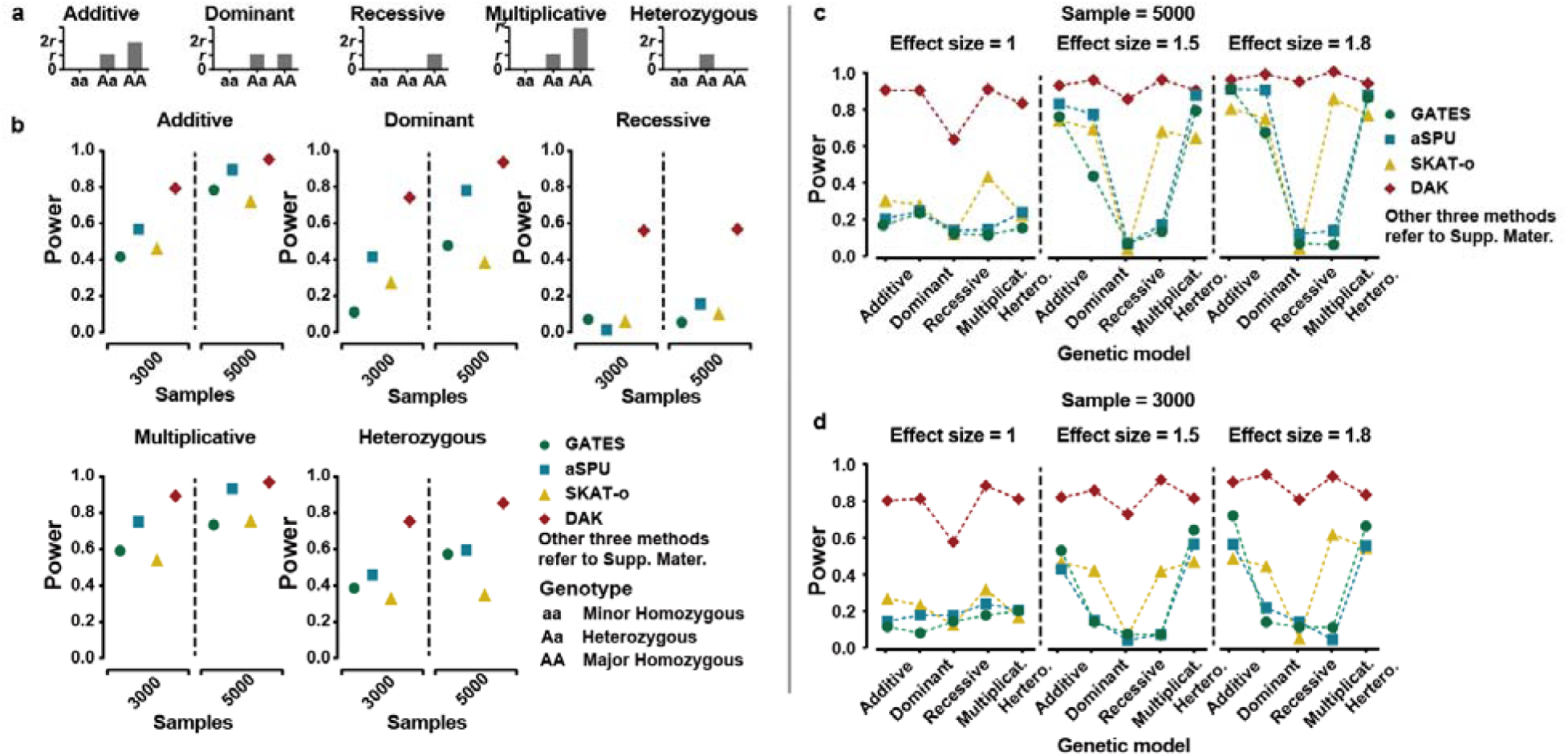
Disease risk levels for different genotypes in five genetic models. (**a**) Performances to discover the disease pathway resulted from single common variant. Effect size was set to 0.2 and simulated phenotypes were generated under five effect models. Under each sample size (3,000, 5,000), seven methods (four showed here and three in **Supplementary Figures**) were used to discover the disease pathway. Power was calculated from 100 replicates after Bonferroni correction. (**b**) Performances to discover the disease pathway resulted from single rare variant. Effect size was set to 1, 1.5 and 1.8, respectively to simulate phenotypes. 5,000 samples were considered. (**c**) Performances of DAK to discover the disease pathway resulted from single rare variant under small sample size (3,000).

**Figure 3.**
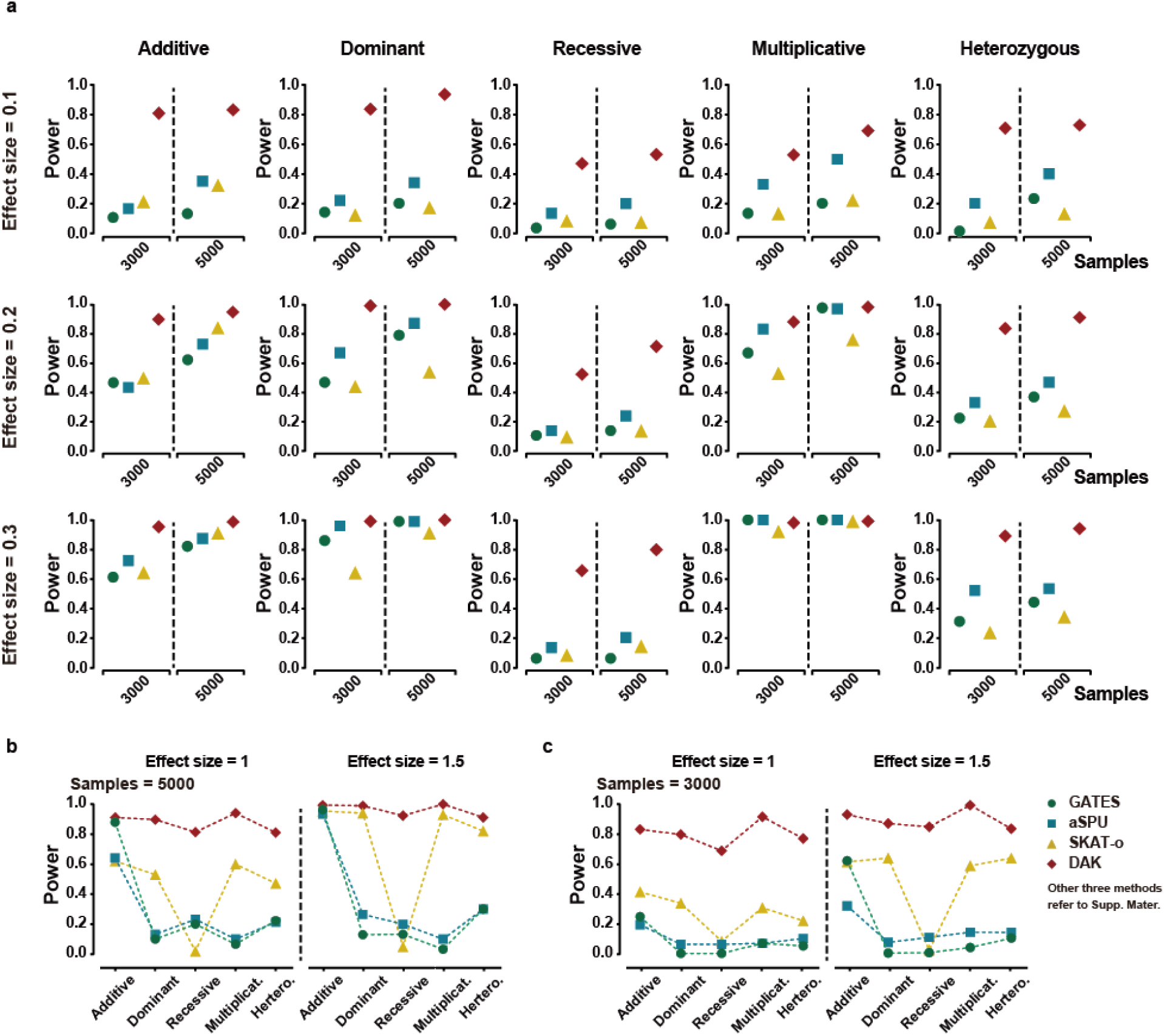
(**a**) Performances to discover the disease pathway resulted from three common variant. Effect size was set to 0.1, 0.2, 0.3 and simulated phenotypes were generated under five effect models. Under each sample size (3,000, 5,000), seven methods (four illustrated here) were used to discover the disease pathway. The power was calculated from 100 repeats after Bonferroni correction. (**b**) Performances to discover the disease pathway resulted from three rare variant. Effect size was set to 1, 1.5 and only 5,000 samples were considered. (**c**) Performances of DAK to discover the disease pathway resulted from three rare variant when only a small sample size (3,000) is available.

### Type-I error

In each simulation experiment, we simulated dataset under null (no causal pathway) or alternative (disease was caused by different genetic associations) hypothesis (**Fig. 2a** and **Methods**). All seven methods were tested on simulated datasets. Performances of different approaches were evaluated using type I error rates (corresponding to null hypothesis) and empirical powers (corresponding to alternative hypothesis) (**Methods**) in 100 replicates.

We first report the Type-I error. If no causal loci existed in all pathways (null hypothesis), all methods showed low error rate level (**Supplementary Fig. 5**). Changing the sample size had little effects on the results. The training curve showed DAK converged within several iterations (**Supplementary Fig. 6**).

### Single effect

We then considered that the disease was caused by a single common variant. To illustrate different functional pathway of genes to the disease, we assumed the allele of the causal locus contributed to the disease in five different genetic models: 1) additive model, minor homozygous genotype had two-fold effect than the heterozygous type; 2) dominant mode, two genotypes showed the same effect size; 3) multiplicative model, minor alleles increased the disease risk exponentially; 4) recessive model, only minor homozygous genotypes had effects; and 5) heterozygous model, only heterozygous alleles had effects (**Fig. 2a**).

On the most widely-used additive disease mode, we found that all methods showed reasonable accuracy to identify the pathway with disease locus (**Fig. 2b** and **Supplementary Fig. 7**). However, when the fundamental genetic model changes, the power of all comparing methods dropped dramatically while DAK maintained reliable performances with best power across all conditions. Specifically, for the challenging recessive genetic model, accuracies of all comparing methods greatly decreased and were far below the performances of DAK. The performance of DAK was further improved when increasing the effect size while other methods were still of low accuracy (**Supplementary Fig. 8**). We further noted that when the sample size was increased to 5,000, the power of all methods were increased and DAK was still the best (**Fig. 2b** and **Supplementary Fig. 7**).

The discovery of rare variants (minor allele frequency < 0.5%) is a challenging task in GWAS due to the low gene frequency. We simulated a rare dataset of 5,000 samples where the disease was caused by single rare variant under five genotype models. Again, DAK obtained much higher performances than others on recessive and multiplicative genetic models (**Fig. 2c** and **Supplementary Fig. 9**). When the effect size was decreased, other comparing approaches failed but DKA can still maintain very reliable performances (**Supplementary Fig. 9**). We demonstrated DAK could discover the causal rare variant at power around 0.8 on datasets even only with 3,000 samples (**Fig. 2d** and **Supplementary Fig. 9**), which was a challenging task for other methods.

### Joint effect

Most diseases are results of the joint-effect of multiple genes. However, it can be more challenging to identify the combined and mixed effect signals from multiple causal variants. Here, we simulated joint-effects by randomly assigning 3 causal common variants and generated phenotype under 5 genetic models (**Methods**). Performances of all methods were much lower compared with results under single variant. However, DAK still dramatically outperformed other methods and achieved the most stable performance among all experiments (**Fig. 3a** and **Supplementary Fig. 10**). The performances of all methods was enhanced when the effect size was increased. The advantages of DAK were more obvious when the causal positions were rare variants. (**Figs. 3b, c** and **Supplementary Fig. 11**)

### Applications to real datasets

We performed DAK on four disease datasets: gastric cancer, colorectal cancer, lung cancer and schizophrenia (**Supplementary Table 1**). After the quality control steps, we divided all SNPs into pathway groups by their genetic coordinates (**Methods**). DAK was optimized on one-hot coded pathways and score test was conducted on each pathway using learned neural network parameters to get the statistical P-value.

For the gastric cancer (GC) dataset, three KEGG pathways exhibited genome-wide significance after Bonferroni correction (*α* = 0.05/186 = 2.68E-4). Two of them (*Terpenoid backbone biosynthesis* and *oxidative phosphorylation*) showed strong associations (**Fig. 4a** and **Supplementary Table 2**). In a previous study, *terpenoid backbone biosynthesis* was identified as a strong relation to Hepatocellular carcinoma (HCC) using miRNA and mRNA high-throughput sequencing^17^. *Oxidative phosphorylation* is closely related to the biological process in mitochondria and it plays an essential role in the development of tumors ^18^. Existing studies have shown its association to endometrial carcinoma, leukemias, lymphomas, etc ^19^. Recent work also indicated it could be an important target to treat cancer using relevant inhibitor ^20^. For the *focal adhesion* pathway, it is important for cell proliferation, cell survival and cell migration. In cancer, activities of focal adhesion were altered during tumor formation and developing^21^. It is also a widely known target for cancer therapy development^22^. For the other three pathways showing borderline significances, *alpha linolenic acid metabolism* was discovered to downregulated human and mouse colon cancers^23^; Function of *ubiquitin mediated proteolysis* on cancers was also widely known^24^.

**Figure 4.**
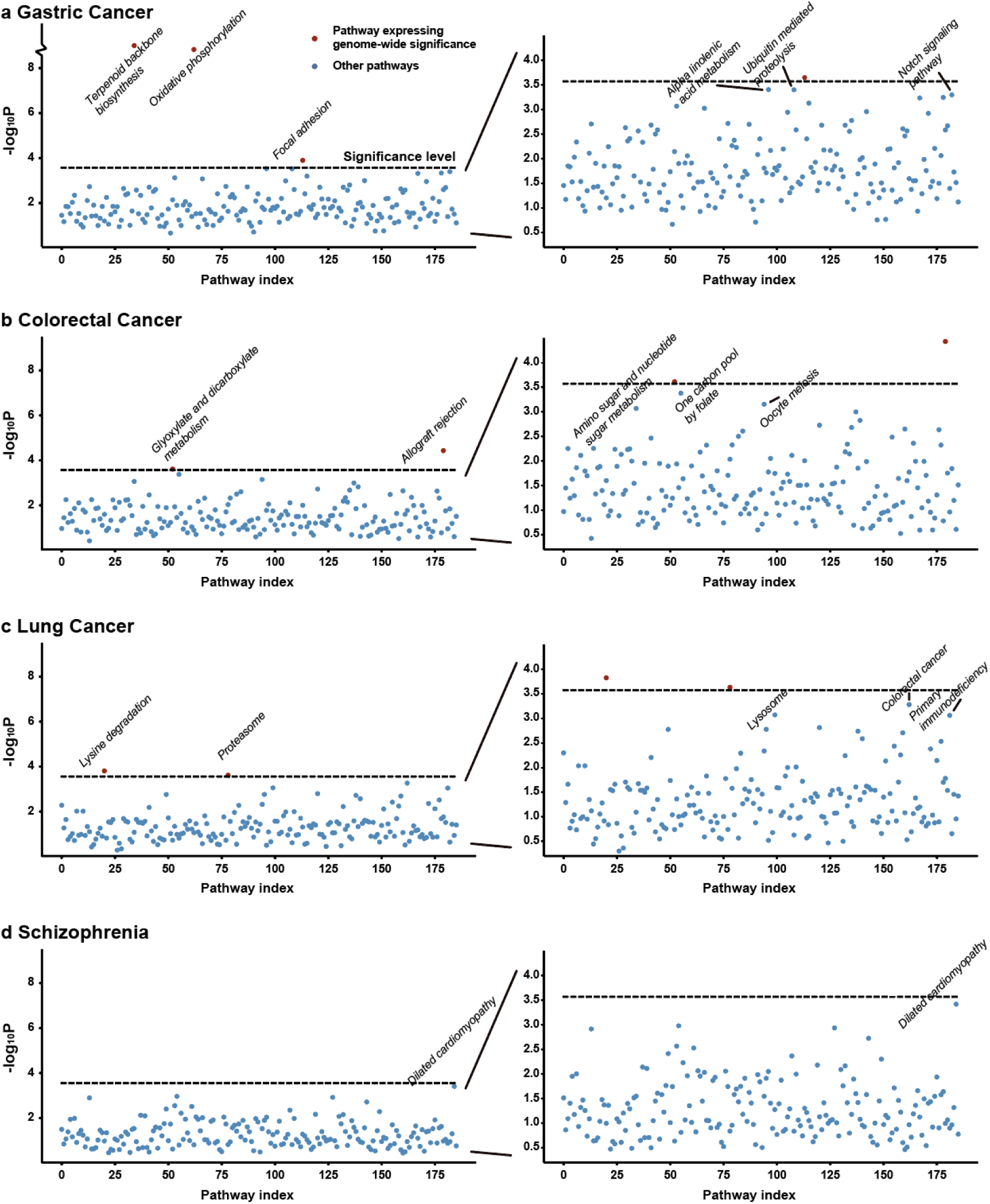
Scatter plots of P-values of KEGG pathways by DAK on four real datasets. Pathways showing genome-wide significances after Bonferroni correction (= 0.05/186 = 2.68E-4) were marked in red.

For the colorectal cancer (CRC) dataset, DAK identified two KEGG pathways showing genome-wide significance (**Fig. 4b** and **Supplementary Table 3**). The most significant pathway, *allograft rejection*, is well known as an immune action pathway. The relation between allograft rejection, blood transfusion and colorectal cancer recurrence was reported at early time^25^. The other significant pathway *glyoxylate and dicarboxylate metabolism* was recently identified to be related to the metabolic switch in colorectal cancer cells ^26^. Other three pathways, *one carbon pool by folate, oocyte meiosis* and *amino sugar and nucleotide sugar metabolism* were also discovered as high risky pathways to CRC. The mechanism between one-carbon metabolism and CRC has been studied^27^ and several key mutations in this pathway has been related to CRC^28^. Oocyte meiosis was identified to be associated to colonic diseases in previous study based on expression data^29^ and amino sugar and nucleotide sugar metabolism may contribute to the lipid metabolism abnormality in CRC^30^.

For the lung cancer (LC) dataset, DAK reported two significant pathways: *lysine degradation* and *proteasome* (**Fig. 4c** and **Supplementary Table 4**). In LC treatment, proteasome inhibitor has been used to non-small cell and small cell LC^31-33^ while lysine modification was discovered to impact a wide range of cancer types^34^. Other three pathways also had relatively small P-values. Colorectal cancer pathway indicates that LC may share causal genes with certain types of CRC. Lysosome was reported to support the development LC^35^. For primary immunodeficiency pathway, it is known to lead to infections and cancers^36^.

For the schizophrenia (SP) dataset, we did not identify pathways reaching genome-wide significance after statistical correction (**Fig. 4d** and **Supplementary Table 5**). Interestingly, one pathway, *dilated cardiomyopathy* (DCM), showed borderline significance with SP. This pathway is related to the heart muscle disease and can lead to heart failure. There is no existing study indicating its biological connection to SP. However, one clinical investigation has shown that after neuroleptics of SP, patients had a significantly increased possibility to get DCM^37^. In other detailed case reports, the usage of clozapine as the treatment to SP finally lead to DCM ^38-40^. This implies that SP and DCM may share biological pathways and the treatment may target at the process that is important to both.

Taken together, DAK efficiently discovered pathways that were known to be associated with diseases and also revealed new potential causal pathways.

## Methods

### DAK architecture

For the *i*-th individual from a total number of *N* samples, *y*_*i*_ denotes the phenotype (such as disease or control); *x*_*i*_ ∈ ℝ^*K*^ is an adjusted vector composed of *K* environmental related factors (e.g. gender, stratification and bias). The genotype of each SNP belongs to one of three types: major homozygous, heterozygous and minor homozygous genotypes. Therefore, it is natural to represent the genotype of each SNP by a one-hot vector with the non-zero entry indicating its particular genotype.

We grouped all *l*^(*p*)^ SNPs on the *p*-th pathway of individual *i* together and get the corresponding pathway-level genotype matrix 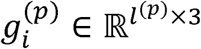 After pathway assembling, we get a total number of *P* pathways for all samples.

We transform each 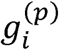 through convolutional layers *conv*(· |Θ_*c*_) with *M* convolutional operators:

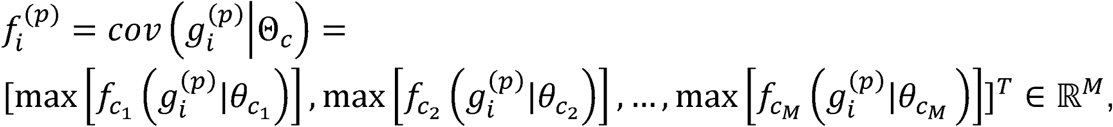

where 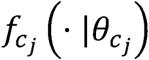 represents the *j*-th convolutional operator with parameter 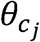 and max [·] is the max-pooling operator. 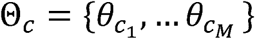 denotes all learnable parameters of the convolutional layer.

By applying the output of the convolutional layers through a *h*_∞_ layer^41^, we obtained the kernel representation of the *p*-th pathway for individual *i*,

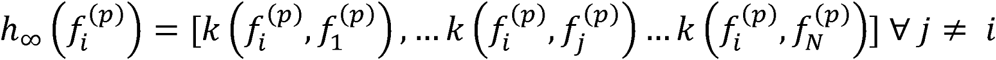

where *k*(·, ·) is a kernel function^12^ and *N* is the number of samples. We then define a pathway-level kernel regression function:

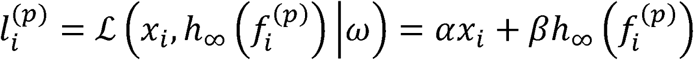

where *ω*= {*α, β*} contains learnable regression coefficients for environment factor and genotype features, respectively. For individual *i*, we can get 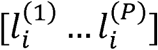 from a total number of *P* pathways.

We noticed that the labels (disease *v*.*s*. non-disease) are only provided at the individual level while not at each single pathway level. We hence consider multiple instance learning loss and define the individual level label for sample *i* as:

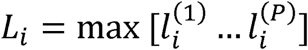

This multiple instance learning loss is naturally explained in the context of GWAS: a sample is treated as a patient if at least one of his pathways is associated with the disease. The training loss is defined as:

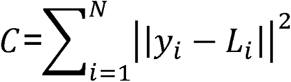

This loss function is optimized by TensorFlow in batches.

After well training, we performed score test to quantify the statistical significance of each pathway using the same approach in SKAT^12^. For each pathway *p*, the statistic score was derived from the kernel similarity matrix 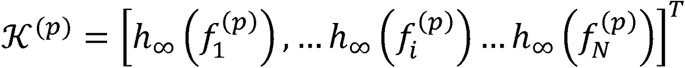 via:

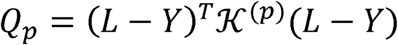

where 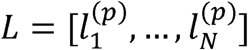(*resp*. *Y* = [*y*_1_, …, *y*_*N*_]) is the predicted (*resp*. ground truth) disease statues for the pathway *p* across *N* samples. As introduced in SKAT, the *Q*_*p*_ was compared with the mixture of *χ*^2^ distributions to obtain P-value.

### Simulation of genotype and data preprocessing

We downloaded haplotypes of CEU population from 1000 Genomes Project^42^. Based on this reference, we simulated full genome data of 10,000 samples using HapGen 2 software^43^. On simulated dataset, we performed the following data quality control steps using Plink^4^: removing individuals with missingness > 0.05; removing SNPs with missing rate > 0.05 or Hardy-Weinberg equilibrium <1e-5. After that, all data were converted into raw files.

### Simulation of phenotype

Phenotypes for samples were simulated based on statistical hypothesis. Under null hypothesis that no causal pathway existed, case / control (represented in 1 / 0) labels were assigned randomly. Under alternative hypotheses, phenotypes were generated using linear models:

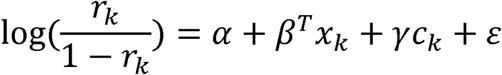

where *r*_*k*_ is the probability for sample *k* being a disease; *x*_*k*_ ∈ ℝ^*K*^ is the vector of environmental factors as mentioned before and *β* ∈ ℝ^*K*^ is the corresponding effect weights; *c*_*k*_ ∈ ℝ is the genotype of pre-selected causal SNP and is coded according to the genetic model assumption. For example, *c*_*k*_ = 0, 1, 0 for the genotype “AA”, “Aa”, “aa”, respectively. For multiplicative genetic model where the disease increased exponentially, we first determine the risk *r*_*k*_ for samples with “Aa” allele and then exponentially increase the risk for “aa” samples. *γ* is the effect size of genotype. We followed the same setting in SKAT^13^, with a 0.2 effect size equivalent to odd ratio of 1.22.

For simulation of disease caused by joint effects, we extend the linear model to

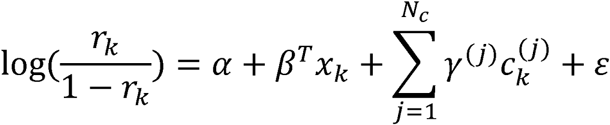

where *N*_*c*_ is the number of causal SNPs. After simulating phenotypes, we randomly selected 50% cases and 50% controls for analyses.

### Pathway set assembling

A total of 186 Kyoto Encyclopedia of Genes and Genomes (KEGG) pathways were downloaded from the database of The Molecular Signatures Database (MSigDB) in the items of “C2: curated gene sets”^44^. The whole-genome SNPs were firstly mapped to genes based on their positions (RefSeq hg19^45^). Then genes grouped in the same pathway were further assembled together.

### Real dataset collections

The genotyping data of GWASs of gastric cancer and schizophrenia were deposited in the database of Genotypes and Phenotypes (dbGaP; phs000361 and phs000021, separately). The genotyping data of GWASs of colorectal cancer and lung cancer were derived from previous studies^46, 47^.

All GWAS datasets were firstly imputed using SHAPEIT and IMPUTE2 based on the 1000 Genomes Project (Phase I, version 3, 1092 individuals. Then the imputed SNPs were cleaned with the criteria of (i) MAF < 0.01; (ii) call rate < 95%; (iii) Hardy–Weinberg equilibrium *P* < 1.0 × 10^−6^; (V) info score < 0.3. The population structure was estimated by a PCA using EIGENSOFT 5.0.1, and the principle components were extracted as covariates, corresponding with age, sex and variables if appropriate for modeling adjustment.

### Evaluation

Performances of all methods were quantified under two metrics: type I error rate and empirical power, corresponding to experiments conducted under assumptions that no disease existed or causal pathway existed. On simulated datasets, all comparing methods were used to derive pathway-level P-values. Under each experimental setting, the association analysis was repeated 100 times on different datasets that were randomly sampled from simulated data. Then, the type I error rate / empirical power was defined as the proportion of experiments detecting significant pathways among 100 repeats.

### Comparison methods

HYST: HYST combines extended Simes’ test and scaled *χ*^2^ test from single SNP association results.

Burden: Burden test uses MAF as weights and additively combines all SNPs.

GATES: GATES takes extended Simes’ test to aggregate single SNP test results.

SKAT: SKAT employs kernels to model the similarity between individuals and directly calculates the association significance between sample kernels and sample phenotypes. Here we used the default kernel setting (“linear.weighted”) and default parameters.

aSPU: aSPU is a method for adaptive test of association analysis. It employs the sum of powered score test to combine single SNPs.

SKAT-o: SKAT-o combines SKAT and Burden test and selects best results from them. We also used the default settings for SKAT.

DAK: The detail structure of DAK was illustrated in **Supplementary Fig. 1**. We also employed linear kernel to be comparable with SKAT. The model was constructed in TensorFlow framework and was run on machine with Nvidia Titan X GPU. We set the training epoch to 100.

### Software availability

DAK is available from Github: https://github.com/fbaothu/DAK

Other tools used in this work can be downloaded from:

Plink: http://zzz.bwh.harvard.edu/plink/

HAPGEN 2: https://mathgen.stats.ox.ac.uk/genetics_software/hapgen/hapgen2.html.

The 1000 Genomes Project: http://www.1000genomes.org/.

UCSC Genome Browser: https://genome.ucsc.edu/

SKAT and SKAT-o: https://www.hsph.harvard.edu/skat/

GATES, HYST and aSPU: https://cran.r-project.org/web/packages/aSPU/index.html

## Acknowledgement

We thank Prof. Xihong Lin from Harvard T.H. Chan School of Public Health for the valuable discussion.

