## Supplementary material for "Explaining the genetic causality for complex diseases *via* deep association kernel learning": Supplemtary Figure

### Supplementary Materials

#### Supplementary Figure 1

The detailed structure of convolutional neural networks used in DAK. Dimensions of layer weights and data were shown.

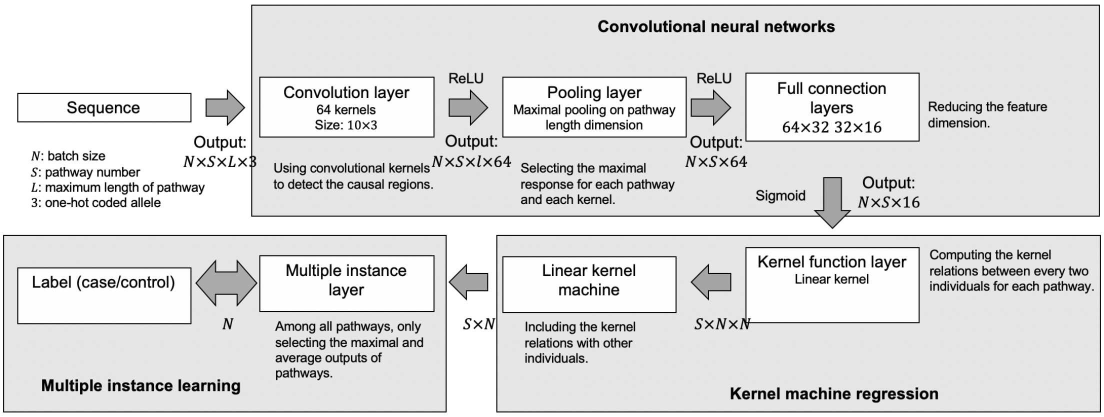

**Supplementary Figure 2**

Kernel machine regression module in DAK. We employed the same framework to conduct gene-based association analysis following the widely used sequence kernel association test (SKAT). For each pathway, deep features were used to construct the kernel similarity matrix by comparing every pair of samples. Then kernel was regressed into logit to indicate the probability of being a case for each sample.

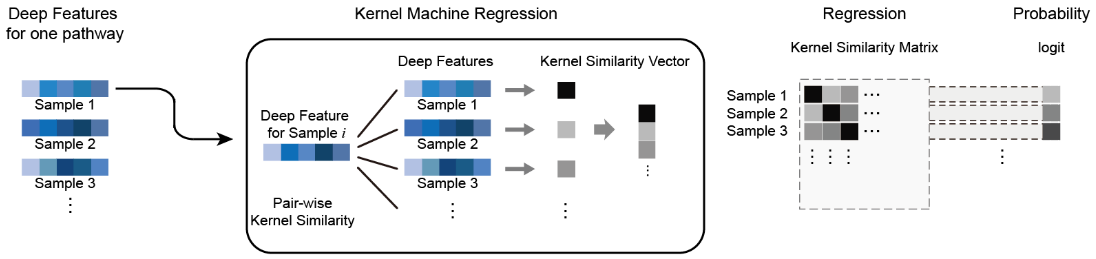

**Supplementary Figure 3**

The association test step of DAK. After training, samples for test were input into the trained model to calculate the kernel matrix in deep latent space for each pathway. Kernel matrices were compared with sample labels and association significances were determined by score test.

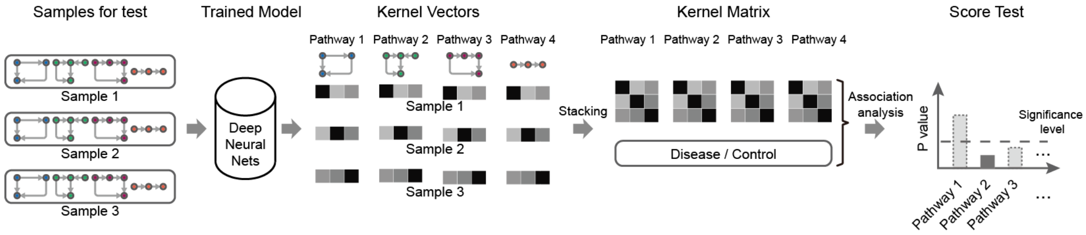

**Supplementary Figure 4**

Model training and significance test stages of DAK. In the training stage, parameters are optimized. In the testing stage, sequence is transformed into deep features and used for significance test.

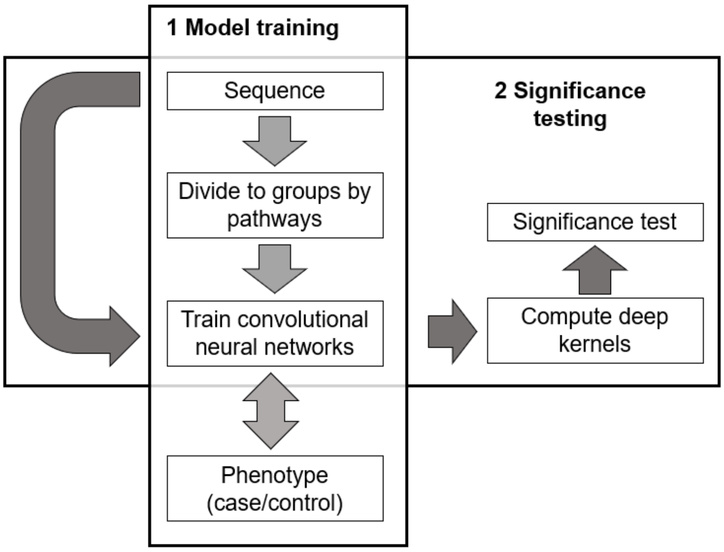

**Supplementary Fig. 5**

Evaluation of type I error rates for all comparing methods. On simulated dataset, phenotypes were generated randomly and each method was used to calculate P-values for all 200 pathways. Type I error rates obtained on 100 repeats (n=100).

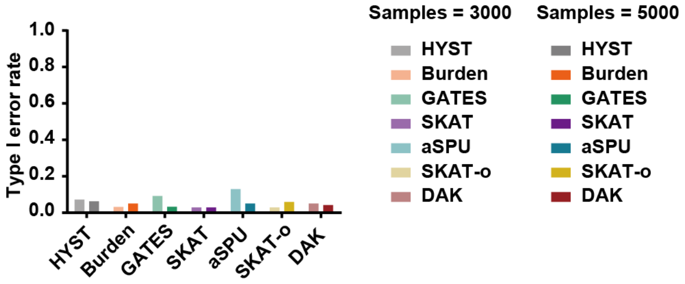

**Supplementary Figure 6**

P-values from DAK association tests using parameters on different training iterations. Average P-values and confidence intervals were obtained on 100 repeats (n=100).

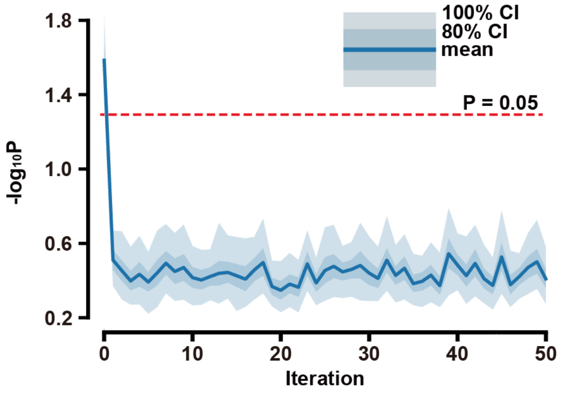

**Supplementary Figure 7**

Performances of all 7 comparing methods on discovering disease pathway. Disease statuses were simulated under effect size of 0.2 (**a**) and 0.3 (**b**) and five effect models were considered. Sample size was set to 3,000 and 5,000, respectively.

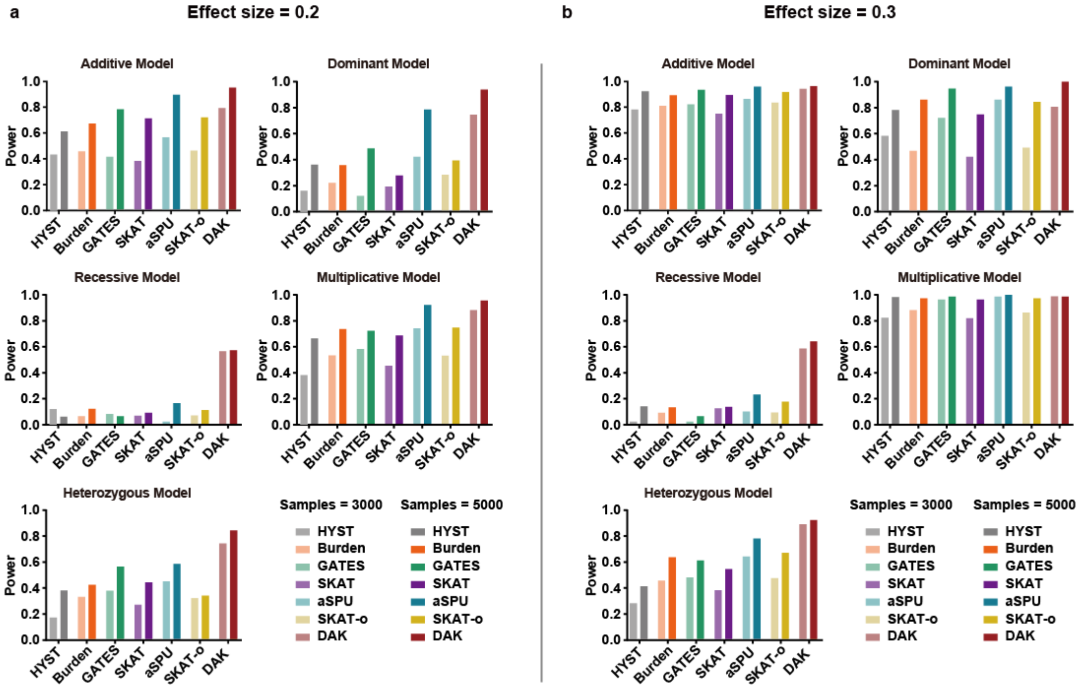

**Supplementary Figure 8**

Powers of all 7 comparing methods on recessive effect model with large effect size (0.4). A single common variant was set as causal locus.

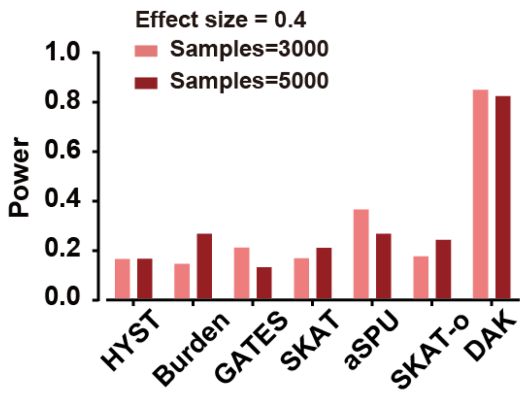

**Supplementary Figure 9**

Powers of all 7 comparing methods on disease caused by single rare variant. Sample size was set to 3,000 (left) and 5,000 (right). Effect size was set to 1, 1.5 and 1.8 to simulate genotypes under five genetic effect models.

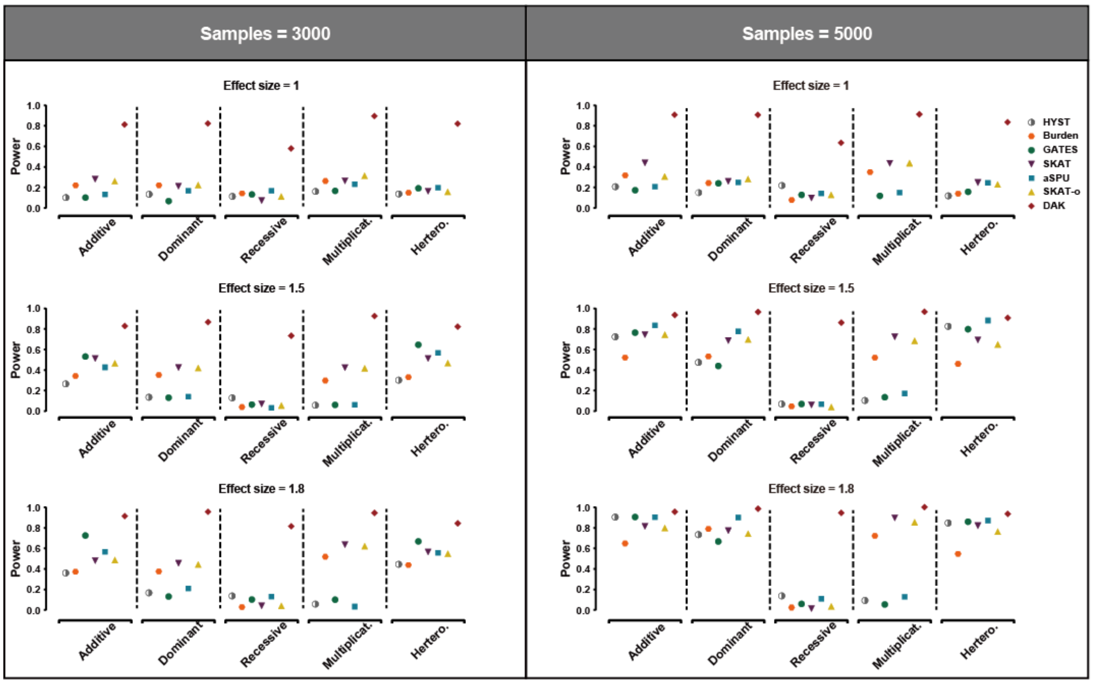

**Supplementary Figure 10**

Performances of all 7 comparing methods on discovering disease pathway resulted by three common variants. Disease statuses were simulated under effect size of 0.1, 0.2 and 0.3 and five effect models were considered. Sample size was set to 3,000 and 5,000, respectively.

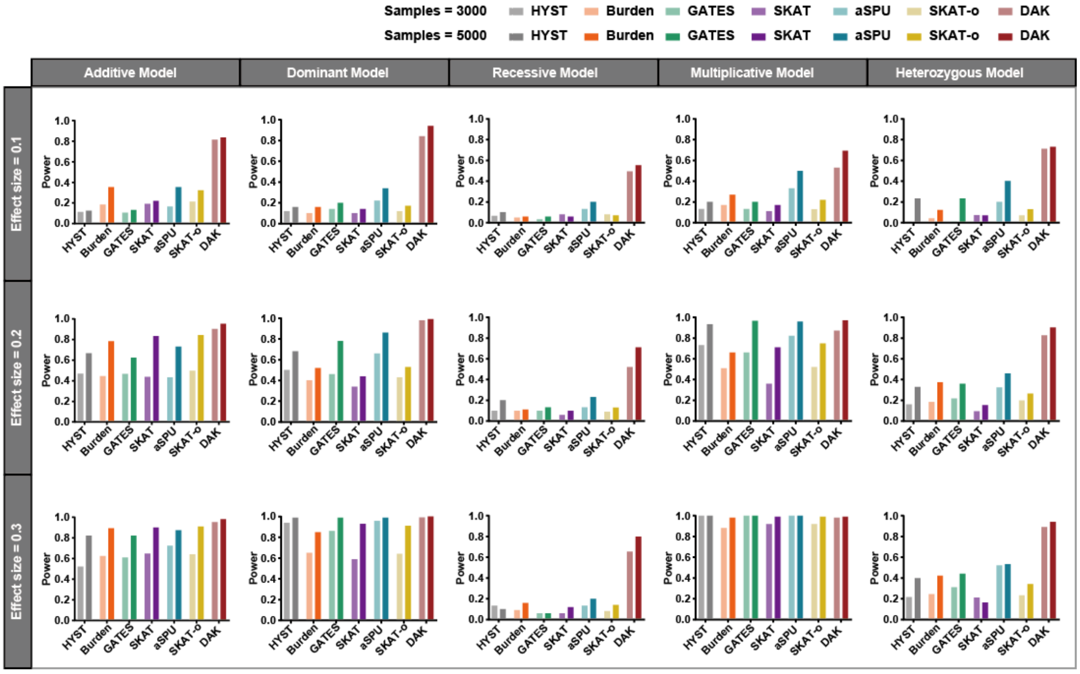

**Supplementary Figure 11**

Performances of all 7 comparing methods on discovering disease pathway resulted by three rare variants. Disease statuses were simulated under effect size of 1 and 1.5 and five effect models were considered. Sample size was set to 3,000 and 5,000, respectively.

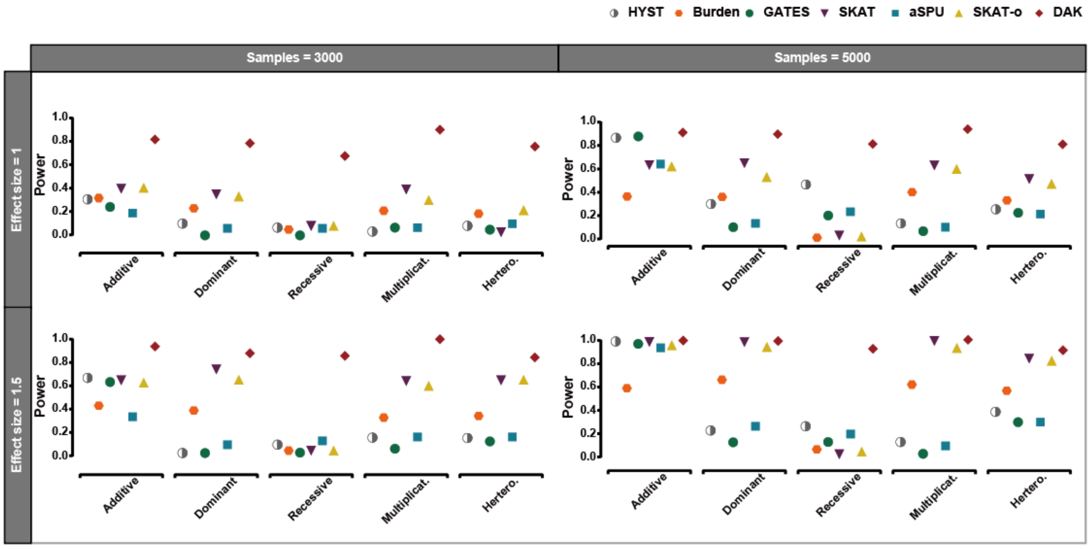

**Supplementary Table 1**

Sample sizes of datasets used in real application.

| | Case | Control |
|---|---|---|
| Gastric cancer | 1625 | 2100 |
| Colorectal cancer | 1316 | 2207 |
| Lung cancer | 984 | 970 |
| Schizophrenia | 1402 | 1442 |

**Supplementary Table 2**

Application of DAK on gastric cancer dataset. Pathways with top ten significances were listed.

| KEGG pathway name | P-values |
|---|---|
| *Terpenoid backbone biosynthesis* | 0 |
| *Oxidative phosphorylation* | 0 |
| *Focal adhesion* | 1.26E-04 |
| *Alpha linolenic acid metabolism* | 2.95E-04 |
| *Ubiquitin mediated proteolysis* | 2.95E-04 |
| *Notch signaling pathway* | 2.98E-04 |
| *Hypertrophic cardiomyopathy hcm* | 4.02E-04 |
| *Systemic lupus erythematosus* | 4.69E-04 |
| *Prostate cancer* | 4.82E-04 |
| *Cell adhesion molecules cams* | 6.42E-04 |

**Supplementary Table 3**
Application of DAK on colorectal cancer dataset. Pathways with top ten significances were listed.

| KEGG pathway name | P-values |
|---|---|
| *Allograft rejection* | 3.71E-05 |
| *Glyoxylate and dicarboxylate metabolism* | 2.43E-04 |
| *One carbon pool by folate* | 4.19E-04 |
| *Oocyte meiosis* | 7.04E-04 |
| *Amino sugar and nucleotide sugar metabolism* | 8.58E-04 |
| *Neurotrophin signaling pathway* | 1.01E-03 |
| *Taste transduction* | 1.50E-03 |
| *Antigen processing and presentation* | 1.88E-03 |
| *Circadian rhythm mammal* | 2.09E-03 |
| *Leishmania infection* | 2.25E-03 |

**Supplementary Table 4**
Application of DAK on lung cancer dataset. Pathways with top ten significances were listed.

| KEGG pathway name | P-values |
|---|---|
| *Lysine degradation* | 1.50E-04 |
| *Proteasome* | 2.33E-04 |
| *Colorectal cancer* | 5.26E-04 |
| *Lysosome* | 8.51E-04 |
| *Primary immunodeficiency* | 8.72E-04 |
| *Antigen processing and presentation* | 1.54E-03 |
| *P53 signaling pathway* | 1.67E-03 |
| *Glycosphingolipid biosynthesis globo series* | 1.68E-03 |
| *Long term depression* | 1.84E-03 |
| *Pathogenic escherichia coli infection* | 1.97E-03 |

**Supplementary Table 5**

Application of DAK on schizophrenia dataset. Pathways with top ten significances were listed.

| KEGG pathway name | P-values |
|---|---|
| *Dilated cardiomyopathy* | 3.81E-04 |
| *Butanoate metabolism* | 1.05E-03 |
| *Hematopoietic cell lineage* | 1.16E-03 |
| *Purine metabolism* | 1.22E-03 |
| *Progesterone mediated oocyte maturation* | 1.88E-03 |
| *Propanoate metabolism* | 2.71E-03 |
| *Porphyrin and chlorophyll metabolism* | 2.97E-03 |
| *Glycosphingolipid biosynthesis globo series* | 3.84E-03 |
| *Dorso ventral axis formation* | 4.31E-03 |
| *Aldosterone regulated sodium reabsorption* | 4.99E-03 |
